# Proteomics on longitudinal serum samples differentiates localized and disseminated Lyme disease

**DOI:** 10.1101/2020.11.18.387993

**Authors:** Benoit Fatou, Kinga K. Smolen, Alexia A. Belperron, Zainab Wurie, Steven H. Kleinstein, Ruth R. Montgomery, Ofer Levy, Linda K. Bockenstedt, Hanno Steen

## Abstract

Lyme disease (LD) is the most prevalent vector-borne disease in North America, with ~300,000 new cases annually in the USA alone. LD is most often recognized by the appearance of the skin lesion erythema migrans (EM) at the tick bite site but also can present with signs of disseminated infection manifesting as multiple EM lesions and/or involvement of the heart, nervous system or joints. In this study, we examined the serum proteome of study participants presenting with either a single EM (localized LD) or early disseminated infection (disseminated LD). Samples collected at the time of diagnosis and at convalescent time points were assessed using our in-house developed MStern blotting-based serum proteomics platform. After technical validation of our platform, the temporal analysis from diagnosis to clinical resolution of the infection demonstrates LD stage-associated pathways activation such as a temporary upregulation of acute phase response specific to the participants with disseminated LD. In addition, we identified the members of the serum amyloid A protein family as potentially promising candidate biomarkers to identify those with disseminated LD. The results of this pilot study demonstrate the feasibility of using our time- and cost-effective sample sparing MStern blotting-based serum proteomics platform to efficiently interrogate proteome changes over time in those suffering infections such as LD. These observations establish a new approach to human serum proteomics, provide fresh insight into the host immune responses associated with disease severity (localized versus disseminated infection) and suggest novel biomarker candidate panels for LD stages.

**Importance:** We investigated the proteome changes of *Borrelia burgdorferi*-infected participants with either a single erythema migrans or early disseminated Lyme disease infection. Using our in-house time-and cost-effective proteomics platform, the temporal analysis from diagnosis to clinical resolution of the infection shows a temporary upregulation of the acute phase response specific to the participants with disseminated infection. Finally, specific protein panels were identified as possible biomarker candidates to categorize those having an initial diagnosis of disseminated manifestation using a reference cohort of acute localized infection and a clinically resolved convalescent phase disease samples from the same Lyme disease participants.

## Introduction

Lyme disease (LD) is caused by the *Ixodes* tick-borne infection with *Borrelia burgdorferi* (*B. burgdorferi*) *sensu lato* spirochetes. Since its emergence in the northeastern United States in the early 1970’s, LD has become the most common vector-borne disease in North America with >300,000 cases per year [1, 2]. Humans are incidental bloodmeal hosts in the life cycle of *Ixodes* ticks and can acquire the infection when *B. burgdorferi* is introduced into the skin during tick feeding. The earliest clinical sign of the disease is the skin lesion erythema migrans (EM) that appears within a month of infection (usually within 7-14 days) at the bite site, often accompanied by a fever, malaise, myalgias and arthralgias [2–4]. Once spirochetes disseminate from this site, clinical signs can be seen elsewhere in the skin, giving rise to multiple EMs, and/or in other organ systems, including the heart (carditis) and nervous system (cranial nerve palsies and meningitis) within the first weeks-months, and in the joints (mono-or pauci-articular arthritis) within a year (average 6 months) post tick bite [5, 6]. Antibiotics are effective at resolving disease and preventing later manifestations, although the time to complete resolution of signs and symptoms may be protracted [7–11]. A minority of patients may experience chronic sequelae, including a proliferative synovitis with autoimmune features (post-antibiotic Lyme arthritis) or persistent fatigue, musculoskeletal pain, and subtle cognitive dysfunction that can be debilitating (post-treatment LD syndrome) [12–16]. The pathogenesis of post-antibiotic Lyme arthritis is thought to reflect a dysregulation of the immune response, whereas the etiology of post-treatment LD syndrome is not well understood [17–21].

Unbiased discovery plasma/serum proteomics is an analytical technique that enables detection and quantification of several hundred proteins without the need for prior knowledge of the blood-based proteins of interest. Blood represents a powerful source for biomarkers as (a) it is easily and readily accessible using minimally invasive collection procedures; and (b) it is the principal medium through which immune-related factors circulate systemically to modulate tissue responses and protect against infection. We recently published a high throughput microtiter plate-based proteomics sample processing method, named MStern blotting, for the fast and efficient processing of neat serum or plasma with all high-abundance proteins remaining in the sample. MStern blotting allows for the rapid processing of 96 individual samples per batch; two such batches are readily processed in a single day [22–25]. The name MStern blotting is derived from western blotting as it uses the same type of hydrophobic membrane for protein retention. Our platform omits the step of depletion of the most abundant proteins without compromising the ability to detect and quantify ~400 classical serum proteins, covering a protein concentration range of up to 7 orders of magnitude. Omitting the widely used depletion provides a more complete view of the immune status as ~40% of the commonly depleted proteins including IgGs, IgAs, IgMs, acute-phase proteins and various components of the complement system have important immunological role [26].

Herein, we employed our MStern blotting-based serum proteomics platform to investigate the serum biomarkers found in individuals with either localized or disseminated LD using samples collected early after the diagnosis and at 1 month and after 3 months during the convalescent period. Mass spectrometry (MS)-based proteomics has been suggested as a key technology in the quest for better blood-based diagnostic biomarkers for LD [5]. We first validated the utility of our platform in the context of LD by comparing our initial findings with the published protein panel generated from a targeted LC/MS proteomics approach to identify candidate biomarkers for early localized LD [5]. Then, we applied our MStern blotting-based serum proteomics platform to the quest of mapping LD changes in the serum proteome over time to identify biomarkers and to better understand the pathophysiology of localized and disseminated LD. We then performed a detailed statistical and bioinformatic time point comparison demonstrating near resolution by the final time point of the proteome differences in those originally presenting with either localized and disseminated LD by the last time point. In the 3 time points, we highlighted the differentially expressed proteins (DEPs) as well as the differentially expressed immunoglobulin domains (DEIgs) for the 2 groups. Next, we conducted a time course analysis from acute infection to disease resolution in those with localized and disseminated LD. Finally, specific protein panels were identified as possible biomarker candidates to categorize those having an initial diagnosis of disseminated LD using a reference cohort of acute localized LD and a clinically resolved convalescent phase disease samples from the same LD participants.

## Results

### Validation of the MStern blotting-based serum proteomics platform using submicroliter sample volumes

We have previously used the MStern blotting-based proteomics platform to map changes in the serum proteome in participants who underwent islet auto transplantation after total pancreatectomy [25]. The main advantages of the platform are its compatibility with small sample volumes (<1 μL of plasma/serum), the capacity for high throughput sample processing (96 individual samples per batch and up to two batches per day even without liquid handling robots), and improved accuracy of peptide and protein quantification due to a reduced number of missing values (**Figure 1**). Our aim in this pilot study was to evaluate the utility of the MStern blotting-based proteomics platform in mapping serum proteome changes to an infectious challenge over time and to determine whether changes could distinguish participants with different disease manifestations. Toward this end, we used serum samples from 12 participants diagnosed with either localized (single EM) or disseminated LD collected at 3 time points encompassing the period of initial presentation through late convalescence several months after diagnosis and treatment when participants had clinically recovered from their illnesses (see **Table 1** and **Supplementary Table 2** for details). The first time point (T1, also defined as Lyme in **Figure 1** and **Supplementary Figure 2**; n = 9) was collected from participants enrolled in the study within 9 days after diagnosis; 7 of the 9 of them had received at least one dose of antibiotics (doxycycline) prior to the blood draw. The second time point blood samples (T2; n = 12) were obtained about one month after the initial diagnosis and in all cases after completion of the initial course of antibiotic therapy. The last time point (T3, also defined as “Resolved” in **Figure 1** and **Supplementary Figure 2**; n = 12) was collected 3 to 4.5 months after the initial diagnosis and corresponds to a return to pre-infection clinical baseline as determined by resolution of the presenting signs and symptoms of the disease.

**Figure 1:**
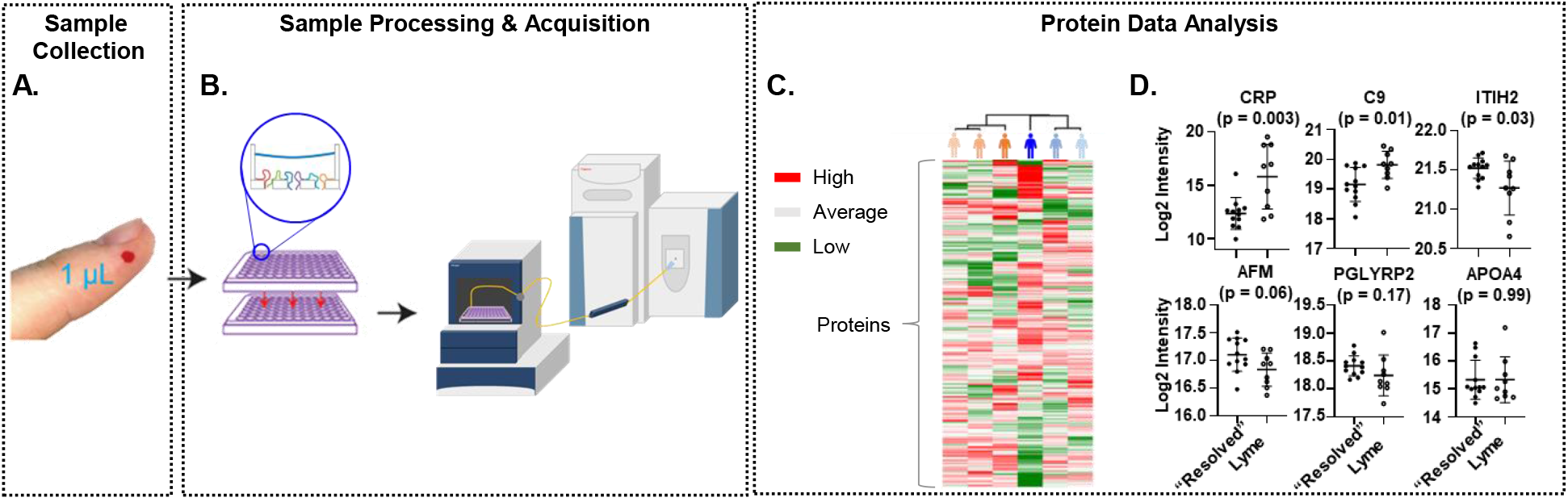
Schematic representation of the MStern blotting-based serum proteomics workflow. (**A**) A starting volume of 1 μl of plasma/serum is required (~50 μg). (**B**) The sample processing part of the MStern blotting-based serum proteomics workflow is based on the use of PVDF microtiter membrane plate and consists of protein denaturation, reduction and alkylation of cysteine residues, rapid protein digestion (2 hours), and purification of peptides. Peptides are separated and analyzed by LC/MS using 45 minutes gradient time (59 minutes run time per sample). (**C**) Data analysis is performed in Spectronaut to obtain protein identification and quantification that could reflect the proteome changes of the stages (localized LD in orange; disseminated LD in blue) of the disease at several time points with a selection of protein candidates based on the study published by Zhou et al [5] (**D**).

**Table 1:** Description of the Lyme disease (LD) cohort

|  | Single EM, Localized LD | Disseminated LD |
| --- | --- | --- |
| Number of participants | 6 | 6 |
| Male | 3 (50%) | 3 (50%) |
| Female | 3 (50%) | 3 (50%) |
| Age (years) | 58.2 ± 17 | 57.5 ± 22.5 |
| Reported Symptoms at Presentation<sup>1</sup> |  |  |
| Fever | 2/5 (40%) | 2/4 (50%) |
| Headache | 4/5 (80%) | 2/4 (50%) |
| Myalgia | 3/5 (60%) | 2/4 (50%) |
| Arthralgia | 2/5 (40%) | 1/4 (25%) |

<sup>1</sup>Due to late study entry at the T2 time point, data for the indicated symptoms were documented at the time of diagnosis (T1) in 5 of 6 localized LD and 4 of 6 disseminated LD participants. T1 Blood samples were collected from 5 of 6 localized LD and 4 of 6 disseminated LD participants. Blood samples were available for all participants at the T2 and T3 time points.

We first analyzed the serum proteome of the above described LD cohort and identified and quantified 8,904 unique peptide spectrum matches (PSMs) from 319 protein groups using a 1% FDR (False Discovery Rate) cut-off. On average 4,632 ± 387 PSMs were detected, enabling identification and quantification of 277 ± 9 protein groups per sample (**Supplementary Figure 1A-B**; **Supplementary Table 1**). These robust numbers confirm the expected reproducibility afforded by the DIA (Data Independent Acquisition) mode. Next, we evaluated the protein dynamic range (**Supplementary Figure 1C**), where the absolute protein concentration was calculated based on the average mass concentrations reported by the Plasma Proteome Database [27]. The concentration range of most of the identified proteins spanned ~6 orders of magnitude from 50 g/L down to 58 μg/L. In addition, we identified 9 proteins with reported concentrations covering an additional 3 orders of magnitude ranging from 6.3 μg/L to 7.3 ng/L. The dynamic range observed in this study was in line with our previous work [25]. Finally, we investigated the data completeness (**Supplementary Figure 1D**). Eighty-seven percent (n = 278) of the detected protein groups were present in at least half of the samples, and 60 % (n = 192) of the detected protein groups were found in all samples. Overall, only 15 % of the protein quantitation values across all samples was missing, which is consistent with published DIA-based proteomics studies [28–31]. In summary, the data quality observed in our study highlights the potential of our MStern blotting-based proteomics platform to identify and quantify a significant number of proteins from non-depleted and unfractionated serum samples. In addition, the low percentage of missing values is key to enable further statistical analysis and interpretation of responses in the study cohort.

To further validate our MStern blotting-based serum proteomics platform in the context of LD, we compared our findings with a published targeted proteomics approach that reported disease-associated abundance differences in blood proteins of LD patients vs. healthy controls [5]. For this verification, we compared T1 and T3 samples, defined as Lyme and “Resolved”, respectively (**Figure 1** and **Supplementary Figure 2**). The T3 time point was considered the most appropriate comparison in our cohort because it is closest to the individual’s immune baseline and these patients reported resolution of LD signs and symptoms [32, 33]. Interestingly, despite the methodological differences with the targeted proteomics study, most (six out of ten) early LD-associated proteins identified in the previous study were also detected in our dataset (**Figure 1**) using an analytical strategy that did not require any prior knowledge. From our study comparison, C-reactive protein (CRP; p = 0.0026), Complement component 9 (C9; p = 0.011), and Inter-Alpha-Trypsin Inhibitor Heavy chain 2 (ITIH2; p = 0.03) demonstrated the most significant differences. Overall, the significant overlap of proteins between the two studies validated our platform and motivated us to further investigate all differentially expressed proteins (DEPs). A more detailed analysis was performed between Lyme and “Resolved” groups enabling identification of potential biomarker candidates to monitor LD infection (**Supplementary Figure 2**).

### Mapping serum proteome changes during recovery from localized vs. disseminated LD

We next continued our in-depth data analysis to assess serum proteome changes characteristic of localized vs disseminated LD and the processes governing convalescence after treatment (**Figure 2**). This figure shows the same set of statistical and bioinformatic analyses comparing localized vs. disseminated LD proteomes in samples collected at each of the three time points (**Figure 2A to C**). First, a principal component analysis (PCA) was performed on all the samples in our study and for clarity, we emphasized the samples from each time point in the respective panels. In addition to the PCA, we investigated the pairwise comparison of disseminated vs. localized LD groups at the global protein level. The relevant volcano plots, bioinformatic analyses of the DEPs using the STRING interaction network, and the number of differentially expressed immunoglobulin domains (DEIgs; p <0.05) are also shown in **Figure 2**. The pairwise comparison at the T1 time point demonstrated the biggest differences between disseminated vs. localized LD groups based on separation along the 2nd principal component in the PCA (p = 0.02) and the highest numbers of DEPs in each group (14 DEPs upregulated in localized LD and 15 DEPs upregulated in disseminated LD) (**Figure 2A**). The significantly changing proteins were further explored using STRING to assess the functional relationships among the DEPs. It was noteworthy that different immune system pathways were activated in disseminated versus localized LD: while the proteins upregulated in localized LD were identified as mostly involved in the complement system, the proteins upregulated in disseminated LD were involved in TLR regulation and hemostasis [33]. The number of DEPs continuously decreased from the T1 to the T2 and the T3 time points, with a similar decrease observed at the transcriptomics level [32, 33]. To our knowledge, this is the first time that LD specific serum proteome changes were identified that could distinguish disseminated vs. localized LD during the course of therapy. The decreasing number of DEIgs in disseminated vs. localized LD (7 vs. 0 at T1, 3 vs. 1 at T2, and 1 vs. 2 at T3, respectively) provides additional evidence for the differences in the humoral immune response detectable in the blood during their respective acute stages.

**Figure 2:**
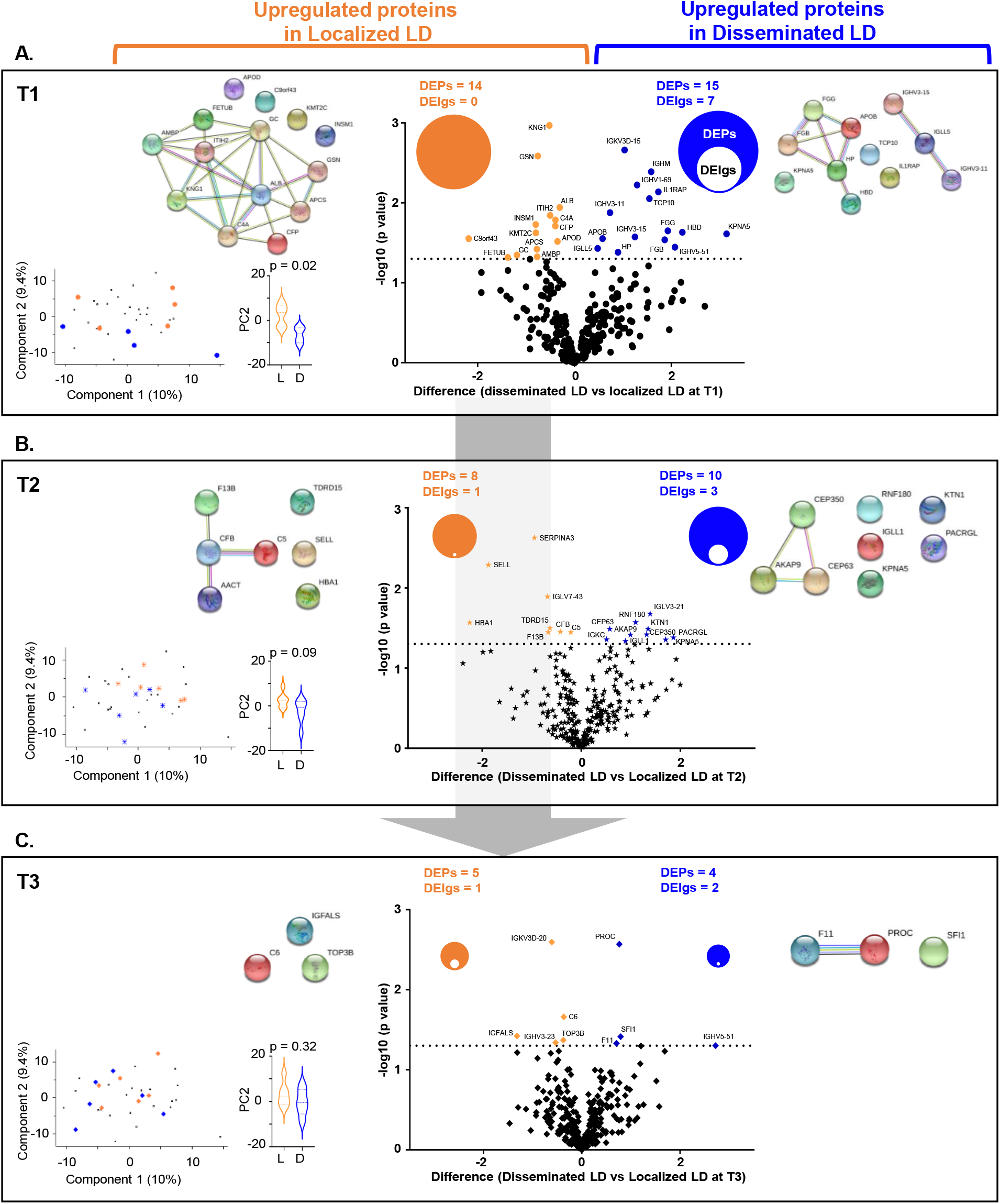
Time point comparison between localized and disseminated LD. Differential analysis between the localized LD (orange) and the disseminated LD (blue) samples at T1 (**A**), T2 (**B**) and T3 (**C**) time points. The number of differentially expressed proteins (DEPs) and immunoglobulins (DEIgs) is also showed.

During LD resolution, the PCA-based separation, the number of DEPs (T2: n = 8 for localized LD and n = 10 for disseminated LD; T3: n = 5 for the localized LD and 4 for disseminated LD), and the number of DEIgs decreased, demonstrating that the localized and disseminated LD groups clearly converged after 3 to 4.5 months, which is consistent with progression to a common physiological state during convalescence [32, 33]. It is noteworthy, that at T2, i.e. 1 month after diagnosis, the different diseases stages were still distinct, as demonstrated by an enrichment of proteins of the complement system, particularly the alternative complement pathway amongst the proteins upregulated in the localized LD group [34].

### Time course analysis from acute infection to disease resolution

Next, we conducted a longitudinal analysis using all three time points in each stage of LD independently (**Figure 3** and **Supplementary Table 3**). We used the STEM (short time-series expression miner) tool to carry out this analysis, and to visualize proteins clusters and trajectories [35], which identified three statistically significant clusters within the protein trajectories of the disseminated LD samples. The co-clustering proteins were subsequently imported into the STRING interaction network to investigate the functional grouping of these relevant clusters. We then selected the same proteins and plotted their trajectories measured for the localized LD samples (**Figure 3**). Very different trajectories of the same sets of proteins observed in either the disseminated LD samples or the localized EM samples are apparent even though in both groups they have resolved to be similar by the last time point. In the first cluster (**Figure 3A**), the protein dynamics for the disseminated LD group decreased from T1 to T2 to remain stable between T2 and T3. Some of these proteins are involved in the acute phase response (highlighted in red) and the complement cascade (highlighted in blue) from the STRING interaction network analysis. In the second cluster (**Figure 3B**), the protein trajectories demonstrate a continuous decrease across the three time points for the disseminated LD and some of these proteins are involved in the negative regulation of proteolysis. Lastly, the third cluster (**Figure 3C**) shows a pronounced increase of protein intensity from T1 to T2 and a stability at T3. Some of these proteins are involved in the regulation of peptidase activity. It is noteworthy that the median trajectories for the same protein set show no change for the localized LD samples. These findings confirm the different proteome profiles observed between localized and disseminated LD from the acute infection to the resolution of signs and symptoms.

**Figure 3:**
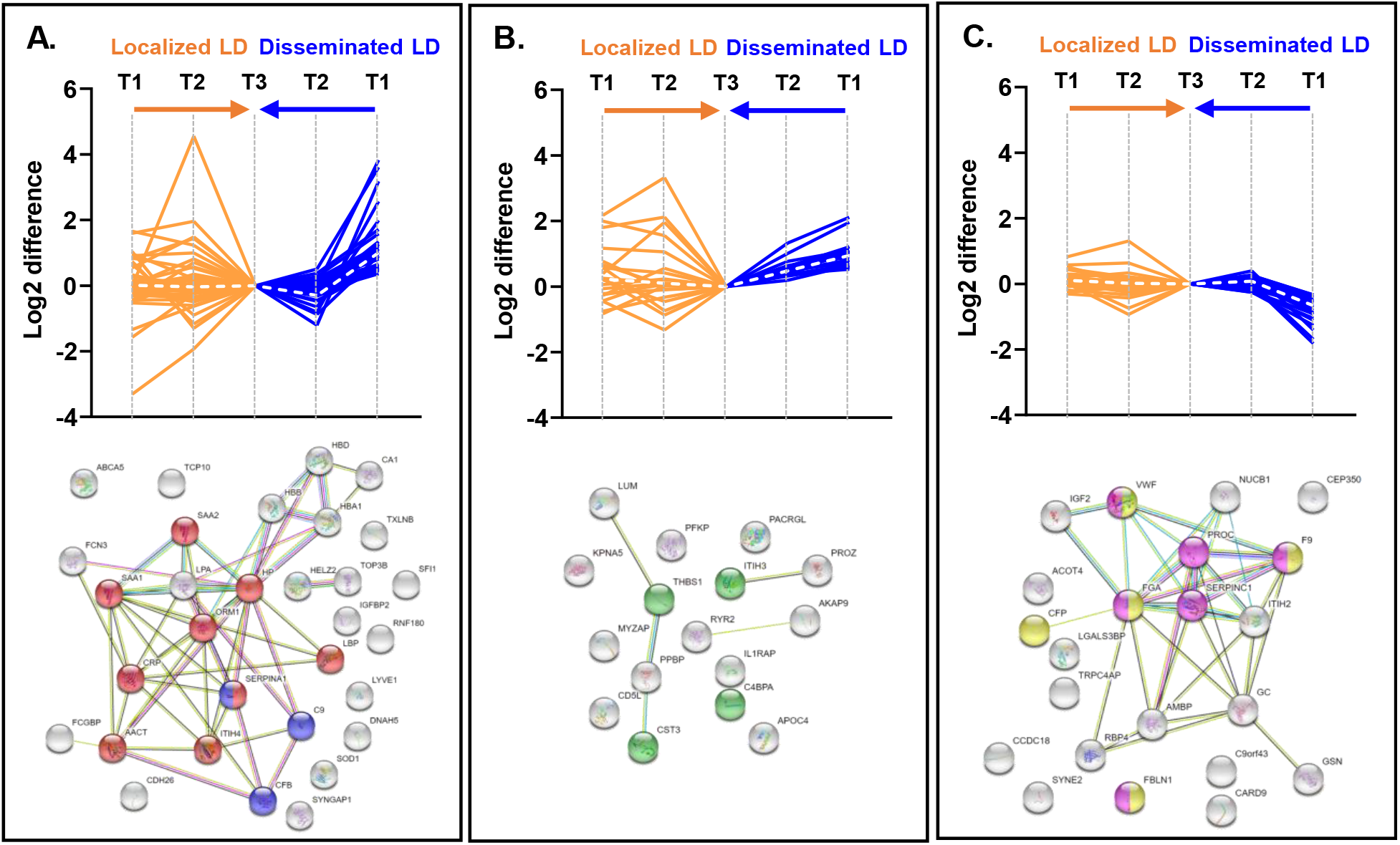
Time course analysis from acute infection to disease resolution in localized and disseminated LD participants. Longitudinal analysis using the STEM program for clustering of the averaged protein intensity profiles across the 3 time points for the disseminated LD group and applying the protein cluster to the localized LD group. For each cluster (**A**), (**B**) and (**C**), the protein names were exported into the STRING interaction network. The white dotted lines represent the median of the Log2 difference of all proteins in the cluster. The proteins highlighted in red are involved in the acute phase response, in blue the proteins involved in the complement cascade, in green the proteins involved in the negative regulation of proteolysis, in yellow the proteins involved in the protein activation cascade, and in pink the proteins involved in the blood coagulation.

### Identifying a biomarker panel for an intended use population mimic

We next sought to model an intended use population to differentiate disseminated LD from non-disseminated LD and other controls. To this end, we created a “control” group comprising the single EM, localized LD group at study entry (T1) and all T3 samples, at which time all participants (both localized and disseminated LD) had clinically resolved infection after treatment. This cohort design ensured that this “control” group also featured patients with acute *B. burgdorferi* infection, allowing us to identify proteins specific to a particular stage of *B. burgdorferi* infection, namely acute disseminated LD, and not only proteins associated with bacterial infection in general. This “control” group was then compared with the disseminated LD samples at T1 (**Figure 4**). We carried out a pairwise comparison, which was visualized using a volcano plot (**Figure 4A**), and a subsequent bioinformatic analysis using the STRING interaction network (**Figure 4B**).

**Figure 4:**
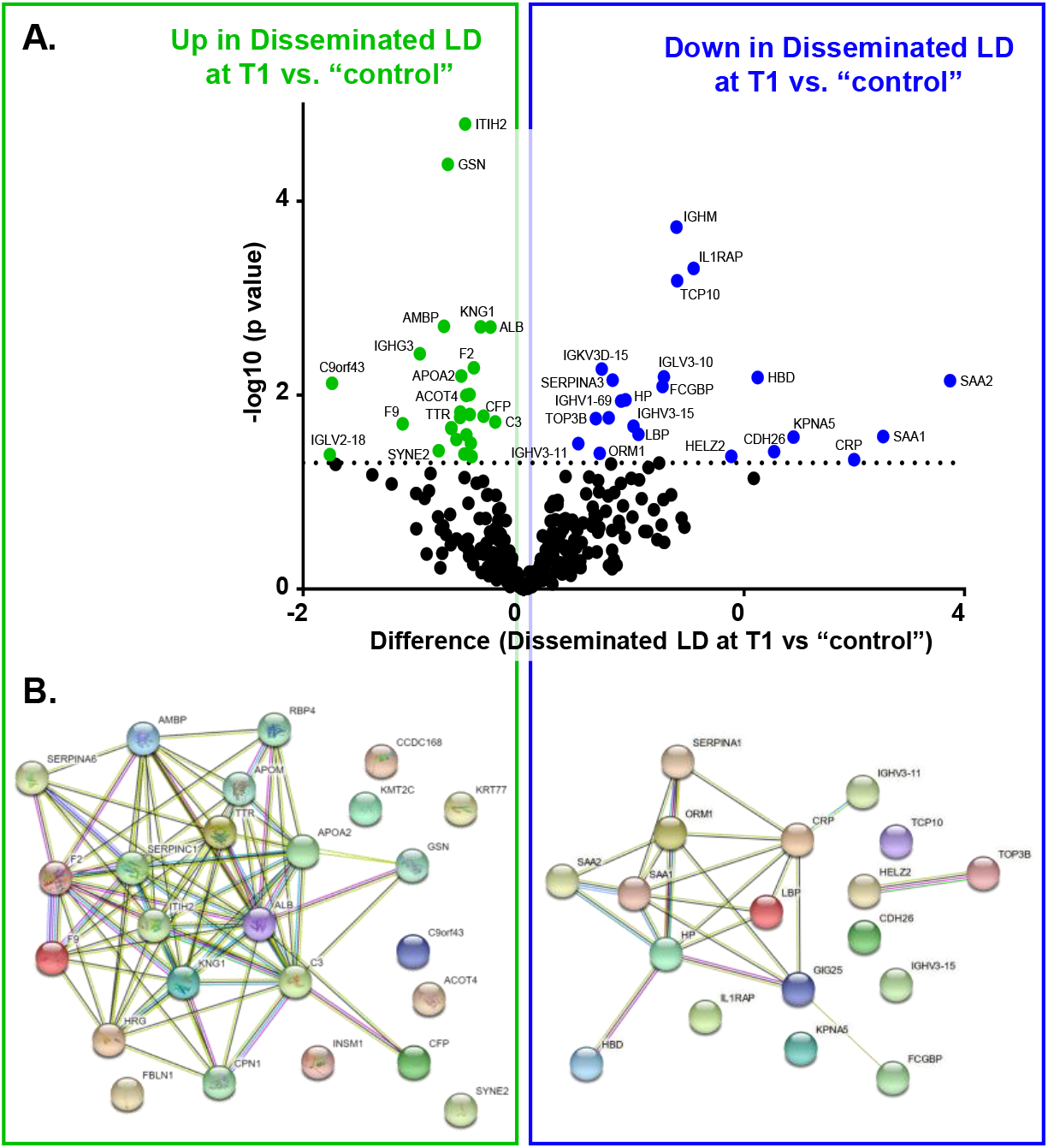
Proteome changes identifies disseminated LD among a “control” group comprised of localized LD T1 samples and localized and disseminated LD convalescent T3 samples. (**A**) Specific protein panel observed in the disseminated LD samples at T1 time point (blue) and the “control” group (green) (all T3 + localized LD T1). (**B**) Protein interaction network of the significant proteins upregulated (blue) and downregulated (green) in the disseminated LD samples at T1 time point.

The pairwise comparison demonstrated a similar number of up- and downregulated proteins (**Figure 4A**). IGHM is the protein that is most significantly upregulated in the disseminated LD group followed by the Interleukin-1 receptor accessory protein (IL1RAP). Both proteins had been observed in previous analyses described above (**Figure 2**). In addition, we observed 10 to 20-fold upregulations for the acute phase proteins SAA1, SAA2, and CRP in the disseminated LD group compared to the “control” group. Using the STRING interaction network (**Figure 4B**), the proteins upregulated in the disseminated LD group were involved in acute-phase, the inflammatory, and the defense responses, highlighting the active state of the innate immune system, which is consistent with the results of the other comparisons described above. The proteins downregulated in the disseminated LD group at T1 were mainly associated with the regulation of proteolysis and peptidase activity. A closer monitoring of the significantly upregulated acute phase response proteins revealed the panel SAA1-SAA2 (AUC = 0.92 ± 0.07; p = 0.011) as a potential candidate biomarkers panel for disseminated LD in an intended use population mimic. In summary, based on the outcome of this analysis, we propose further studies of SAA1, SAA2, CRP, and IL1RAP or combinations thereof as potential biomarkers or biomarker panels for supporting the diagnosis of disseminated LD.

## Discussion

Our study demonstrates the utility of our MStern blotting-based serum proteomics platform in assessing changes in the serum proteome during the course of an infectious challenge. Using serum samples from participants presenting with LD manifesting clinically as localized infection (single EM lesion) versus disseminated infection, we measured the evolution of changes in the serum proteome during acute illness through convalescence and return to a clinical pre-infection baseline. Our unbiased discovery proteomics MStern blotting-based serum platform recapitulated major aspects of a previously published LD plasma proteomics study using a Multiple Reaction Monitoring (MRM)-based LC/MS assay [5]. Our MStern blotting-based proteomics platform has advantages over the platform used in the previous publication as our platform requires ~1% of the starting material (0.3 μL vs 25 μL), utilizes only 50% of the gradient time (45 minutes vs. 90 minutes), and is significantly less labor and cost intensive by omitting steps involved with antibody-based depletion of the highly abundant serum proteins (**Supplementary Table 4**).

The MStern blotting-based proteomics platform applies a DIA-based strategy which combines some of the strengths of targeted methods with those from DDA (Data Dependent Acquisition)-based discovery experiments including the significant decrease in missing values. The reduced sparseness facilitates more robust protein quantification and subsequent statistical analysis [28–31]. Despite the small sample size, our bioinformatic and statistical data analysis allowed us to define features of the proteome and pathophysiological processes that are associated with localized or disseminated LD, and to identify potential biomarker candidates that distinguish these presentations which can be validated in a larger independent cohort.

The availability of longitudinal samples from localized and disseminated LD patients enabled investigation of LD stage-specific biological processes activated during the acute illness and subsequent convalescence. The progressive convergence of our study groups in the PCA across time, including continuously decreasing numbers of DEPs (from 29 to 9) and DEIgs (from 7 to 3) suggest a return to a common post-infectious physiologic state, despite the differences in disease presentation (**Figure 2**). The most striking differences between localized and disseminated LD were the involvement of aspects of the complement and coagulation pathways in localized LD in contrast to the involvement of the pathway associated with the regulation of TLR by endogenous ligands in the disseminated LD stage [36]. Similarly, the pronounced increase in numerous Ig domains in disseminated LD demonstrates the more robust engagement of humoral immunity that is detectable in the serum of patients with disseminated LD. This observation may reflect enhanced B cell response to the pathogen as well as a longer duration of subclinical infection in participants with disseminated LD in comparison to those with single EM, localized LD.

The differences in the convalescence processes for the two investigated LD stages were also underscored by the STEM-based analysis of the temporal protein profiles (**Figure 3**). Only in the disseminated LD stage were protein profile clusters of statistical significance identified. When plotting the proteins profiles of the same protein from the localized LD samples, on average very little, if any, temporal variation was observed.

Our analysis of DEPs identified three proteins of interest that showed localized LD-specific abundance increases in comparison to disseminated LD: ITIH2, kininogen-1 (KNG1), and gelsolin (GSN) (Figure 3A). ITIH2 [37] is one of the biomarker candidates identified by Zhou *et al.* [5]. KNG1 is an inhibitor of thiol proteases and a precursor protein to both high- and low-molecular weight kininogens. High molecular weight kininogen is a cofactor in the human contact system that contributes to the acute inflammatory response and host defense against pathogens [38]. Its cleavage results in the release of bradykinin and antimicrobial peptides, and reduction in its blood levels has been associated with bacterial sepsis [39]. GSN is a multifunctional protein important for intracellular actin multimerization and cytoskeletal remodeling [40]. The secreted form of GSN acts as an extracellular actin scavenger in blood and also modulates immune responses to bacteria through its binding of bacterial cell wall components and reduction of TLR-mediated inflammatory processes [41, 42]. GSN concentrations in blood are diminished in bacterial sepsis [43, 44] and in tick-borne encephalitis and Lyme neuroborreliosis, possibly due to GSN binding of actin released from dying cells and sequestration of inflammatory mediators [41]. The greater abundance of KNG1, ITIH2 and GSN in those presenting with localized LD in comparison to those with disseminated LD suggests that, despite the appearance of a localized infection, systemic changes are present in the blood.

Given this observation of LD stage-associated processes, we searched for specific biomarkers of disseminated LD. Of note, diagnosing LD manifestations that include one or more EM lesions is of lower importance as the skin lesion itself is a sensitive ‘biomarker’ for LD with skin involvement. To this end, we assembled two cohorts to mimic the intended use population: i) acutely disseminated LD vs. ii) non-disseminated LD, which included convalescent disseminated LD patients, and acutely ill and convalescent localized LD patients. By including the acutely ill localized LD patients, we minimized the risk of identifying generic bacterial or *B. burgdorferi* infection markers. This analysis resulted in several promising biomarker candidates seemingly specific for disseminated LD among this “control” cohort.

These potential biomarkers include IL1RAP, a protein that belongs to the receptor complex binding IL-1 which then induces the synthesis of acute phase and proinflammatory proteins during an infection [45]. SAA proteins (SAA1 and SAA2) are the most promising biomarker candidates separating disseminated LD in the context of single EM, localized LD (T1) and “Resolved” (T3) samples (**Figure 5**). SAA proteins are acute phase proteins also produced by the liver and exhibit significant immunomodulatory activity. They can induce the synthesis of cytokines, activate the inflammasome cascade, and serve as chemokines for neutrophils and mast cells [46]. Interestingly, these 2 proteins share up to 90% sequence homology and cannot be distinguished by immunoassays as the antibody reacts indistinctly with both SAA1 and SAA2 [47]. In contrast, mass spectrometry-based methods can readily differentiate between these two proteins.

In summary, we demonstrated that the proteome changes related to the stages of LD can be monitored using microliter volumes of serum samples. The DEPs identified and their related biological pathways may improve the understanding of the LD pathogenesis early after the diagnosis, and during the early and late convalescent period. Furthermore, despite our small sample size, we identified biomarker candidates associated with disseminated LD, namely SAA1, SAA2, and IL1RAP or combinations thereof. Validating these proteins as biomarkers should be the objective of a larger well powered follow-up validation study. In addition to such validation of diagnostic biomarkers for disseminated LD, a biomarker panel for monitoring recovery from LD should be investigated as complete resolution of LD symptoms can be lengthy and the causes of lingering symptoms remain undefined. Such biomarker for recovery would require the collection of later time points, i.e. 3, 6 and/or 12 months post initial diagnosis.

## Methods

### Cohort design / patient characteristics

The cohort was obtained with written informed consent under the guidelines of the Human Investigations Committee of Yale University School of Medicine from patients presenting with disease at Yale New Haven Hospital or Mansfield Family Practice in New Haven and Storrs, CT respectively. The study participants (N = 12) diagnosed with LD based on the CDC case definition [2] and were stratified into 2 groups: those with a single erythema migrans (EM) lesion (localized LD; N = 6) and those with clinical signs of disseminated infection (disseminated LD; N = 6). The disseminated LD group included neurologic disease, systemic flu-like illness that was confirmed by seroconversion, carditis and arthritis. Serum samples were collected at 3 different time points: 1) early after diagnosis, range 0-9 days (defined as T1 or Lyme), 2) at convalescence, about 30 days post diagnosis (defined as T2), and 3) up to 4.5 months post diagnosis, range 3-4.5 months (defined as T3 or “Resolved”). Additional details regarding this cohort can be found in **Table 1** **and Supplementary Table 2**.

### 96-well sample processing/digestion and cleanup

Sample processing employed an MStern blotting-based proteomics platform we have developed as described previously [23, 25]. In brief, 1 μL of serum (~50 μg of proteins) was mixed in 100 μL of urea buffer. Following reduction and alkylation of the cysteine side chains, 10-15 μg of proteins were loaded on to a 96-well plate with a polyvinylidene fluoride (PVDF) membrane at the bottom (Millipore-Sigma), which had been previously activated and primed. Trypsinization of the proteins adsorbed to the membrane was achieved by incubation with the protease for 2 hours at 37°C. Resulting tryptic peptides were eluted off the membrane with 40% acetonitrile (ACN)/0.1% formic acid (FA). The peptides were subsequently desalted using a 96-well MACROSPIN C18 plate (TARGA, The NestGroup Inc.).

### DIA (Data-Independent Acquisition) sample acquisition

The samples were analyzed on the same LC/MS system as the data-dependent acquisition (DDA) runs using identical LC parameters (45 minutes gradient, 59 minutes total runtime). The m/z range 375−1200, covering 95% of the identified peptide, was divided into 15 variable windows based on density, and the following parameters were used for the subsequent DIA analysis: resolution 35000 @ m/z 200, AGC target 3e6, maximum IT 120 ms, fixed first mass m/z 200, NCE 27. The DIA scans preceded an MS1 Full scan with identical parameters yielding a total cycle time of 2.4s.

### Spectral library generation and DIA data analysis

We use a previously published in house generated spectral library [25]. All DIA data were directly analyzed in Spectronaut v12.0.20491.18 (Biognosys, Switzerland). Standard search settings were employed, which included enabling dynamic peak detection, automatic precision nonlinear iRT calibration, interference correction, and cross run normalization (total peak area). All results were filtered by a q-value of 0.01 (corresponding to an FDR of 1% on the precursor and protein levels). Otherwise default settings were used.

### Statistics

The protein abundance/samples matrix was exported into Perseus software [48] where a log2 transformation followed by imputing the missing values for each column from a normal distribution (width = 0.3; down shift = 1.8). For better visualization, the results were exported into GraphPad Prism software v8. The area under the ROC curve (AUROC) calculation for the biomarker panels employed IBM’s SPSS software. To minimize between individual variation, we used the last time point (T3) as “Resolved” as patients at this time point have clinically resolved their acute infection after treatment [32, 33]. Given the pilot nature of this study and limited number of samples, no multiple testing corrections were applied. The DEPs were exported into the STRING interaction network to assess their functional relationships [49]. We used the short time-series expression miner (STEM) tool to visualize protein profiles and dynamics among the 3 time points of localized and disseminated LD groups separately [35]. The averaged protein intensities were calculated for each time point and then the data were normalized such that the time point T3 was set to 0.

## Author Contributions

Manuscript was drafted by BF and KKS; RM, SK, LB designed the clinical study; AB and LB were responsible for sample collection, handling and distribution including clinical data capture; BF, HS designed the analytical strategy; BS, KKS, ZW processed and analyzed the samples; BS, KKS, SK, HS analyzed the data. All authors listed above contributed to the data analysis and/or interpretation and reviewed various versions of the manuscript including the final copy as submitted.

## Acknowledgments and Funding

The authors thank the individuals that participated in this study. This manuscript study was supported by an administrative supplement to OL and HS from the NIAID/NIH to the Human Immunology Project Consortium (U19AI118608), and to RH and LB (U19AI089992).

OL’s and HS’ Laboratories are supported in part by US National Institutes of Health (NIH)/National Institute of Allergy and Infectious Diseases (NIAID) Human Immunology Project Consortium (HIPC) award U19AI118608. The *Precision Vaccines Program* acknowledges the support of the Boston Children’s Hospital’s Department of Pediatrics, Chief Scientific Office and the Boston Children’s Hospital Trust.

## Data deposition

All data are publicly available. The mass spectrometry RAW data and search results have been archived on ImmPort (https://immport.niaid.nih.gov/home) under accession numbers SDY1394 and SDY1395.

